# ChassiDex: A microbial database useful for synthetic biology applications

**DOI:** 10.1101/703033

**Authors:** B P Kailash, D Karthik, Mousami Shinde, Nikhita Damaraju, Anantha Barathi Muthukrishnan, Shashi Bala Prasad, Guhan Jayaraman

## Abstract

ChassiDex is an open-source, non-profit online host organism database that houses a repository of molecular, biological and genetic data for model organisms with applications in synthetic biology. The structured user-friendly environment makes it easy to browse information. The database consists of a page for each model organism subdivided into sections such as Growth Characteristics, Strain diversity, Culture sources, Maintenance protocol, Transformation protocol, BioBrick parts and commonly used vectors. With tools such as CUTE built for codon usage table generator, it is also easy to generate and download accurate novel codon tables for unconventional hosts in suitable formats. This database was built as a project for the International Genetically Engineered Machine Competition in 2017 with the mission of making it easy to shift from working with one host organism to another unconventional host organism for any researcher in the field of synthetic biology. The code along with other instructions for the usage of the database and tools are publicly available at the GitHub page. We encourage the synthetic biology community to contribute to the database by adding data for any additional or existing host organism.

https://chassidex.org; https://github.com/ChassiDex

## Introduction

Synthetic biology is the engineering of biology which involves designing new biological systems that don’t exist in nature or developing new functions in an already existing biological system. In this field, the advancements have been mostly achieved through the use of microorganisms.(1) The DNA of various microbes like *Escherichia coli* have been modified extensively to impart new functions and improve focus on specific applications like bioremediation(2), or production of important biofuels.(3) Similarly, projects which involve the design and building of pathways that enhance the production of the required compound or be responsive to certain environments are incorporated into the microorganisms.(4) Hence the use of various micro-organisms has been instrumental in synthetic biology. A few microorganisms, for example *Escherichia coli*, *Lactobacillus lactis*, *Bacillus subtilis*, *Pichia pastoris* are most commonly used, due to a number of reasons. Multiple factors, like the availability, available parts, and tools to work with, ease of handling the microbe, doubling time, optimum temperature, etc dictate the use of these microbes. While these factors play an important part in the design of the experiments, the microorganisms are used regardless of whether they are well suited or best for those experimental conditions. In some cases, the model microorganisms are engineered to do functions that other non-model microorganisms can naturally do.(5) Microbes like *Vibrio natriegens* has a fast doubling time.(6) *Acinetobacter baylyi* is a naturally transformable bacteria.(7) These bacteria have natural abilities that have to be engineered in conventional bacteria. Hence, it is important that the scientific community take note of the unconventional microbes and the importance of their applications in engineering synthetic frameworks.

The major drawback of working with a non-conventional chassis is the unavailability of tools and parts for the microorganisms, and literature among other things. Due to the dearth in the amount of information regarding various unconventional chassis, the importance of these chassis have been lost on us.(5) While some laboratories around the world have started to work on various microorganisms to develop robust and biological chassis, this information is not necessarily freely available.

To combat this difficulty, we propose ChassiDex. ChassiDex was developed with the aim to curate information about various non-conventional chassis. The curated data is obtained from groups and labs working on the non-conventional chassis. We believe this platform will help data, regarding the handling and editing procedures of the non-conventional microorganism, be readily available for any synthetic biologist to access. Through ChassiDex, we aim to ease the process of accessing data about and encourage the scientific community to use non-conventional chassis.

## The Website

ChassiDex is hosted on the domain chassidex.org, using the services of Netlify (www.netlify.com). All the data and code are stored in our GitHub repository. The website is set up for continuous deployment using a custom made python script, which means any changes made on the GitHub repositories are automatically and immediately reflected on the website. The home page contains a list of organisms and supporting tools available on ChassiDex as well as a tag-wise listing of organisms. Each organism has its own page, where its data is displayed.

## Collection and curation of data

As of April 2019, ChassiDex consists of data for 14 species of microbes commonly used in Synthetic Biology. The data consists of growth characteristics, culture sources for growing the microbe, maintenance protocols, transformation protocols, BioBrick™ parts, vectors, metabolic models, and other online resources and databases that are useful for working with the organism. Each chassis is classified into search-able tags (please see Figure 1) based on the biological nature of the organism and nutritional sources.

**Fig. 1.**
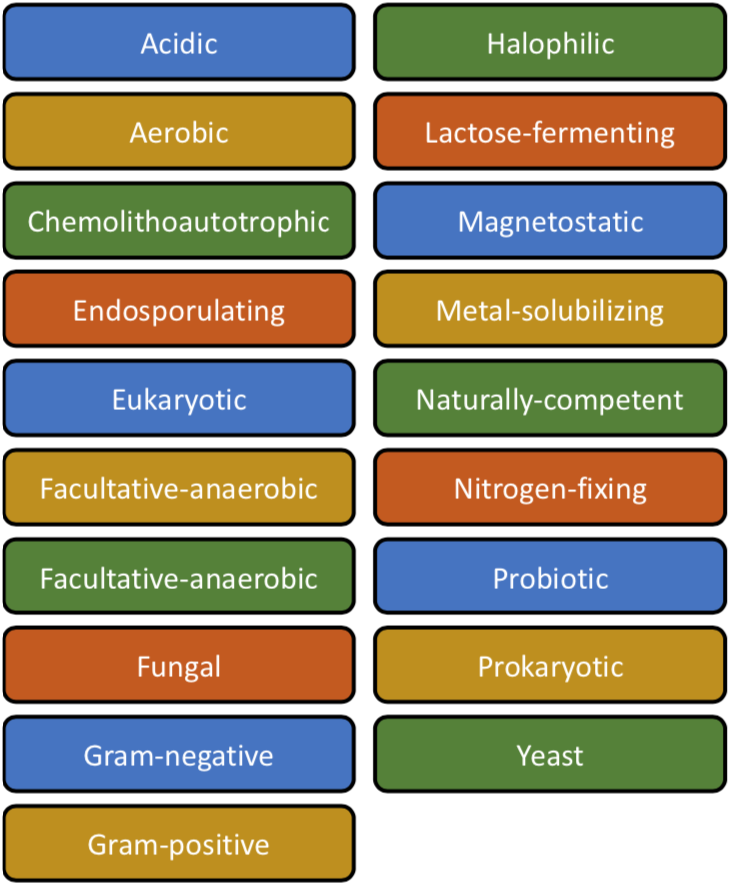
Tags used in categorizing of the data for greater searchability

The collection of data for each chassis begins with a literature survey to find the basic characteristics. This is followed by finding existing relevant data banks for each organism such as EcoliWiki (9) for *Escherichia coli*. An online form (see Supplementary S1 for links) consisting of all the fields in the database is used to enter the data to be sent to the curation team. The data currently hosted on the website has been curated by iGEM 2017 & 2018 team members of Team IIT-Madras. The data for 4 organisms was curated by the respective iGEM teams mentioned as a collaborative effort. (see Supplementary S2)

## Organization of host organism data

ChassiDex uses a flat file for each organism to store data and a custom static site generator script is used to generate the web pages for each organism and the home page. Hence, on landing on the home page, links to the web pages of each organism are provided.

Under the hood, data for each organism is stored in the markdown format. Each of these markdown files has a header with the name of the organism, tags to classify them, and a color code that represents the type of organism it is. The color codes are shown in table 1. The set of markdown files are processed by a python script to generate web pages for each organism. The script also generates the comprehensive and tag-wise listing of organisms on the home page. Various data fields we intend to fill for each organism are provided in table 2. Data fields mostly consist of objective data and we intend to minimize descriptive paragraphs, to move towards better indexing.

**Table 1.** Color coding of organisms in ChassiDex

| Color code | Description |
| --- | --- |
| Red | Gram-negative bacteria |
| Blue | Gram-positive bacteria |
| Purple | Fungi |
| Green | Others |

**Table 2.** A non-exhaustive list of data fields covered for organisms in ChassiDex

| Field | Description |
| --- | --- |
| Applications | Use cases for the organism |
| Strains | Different strains of the organism and their uses |
| Growth Cycle | Growth cycle describing doubling time, etc. |
| Culture suppliers | Stock collections and labs that one can procure the microbe from |
| Growth Media | Media and conditions for cultivating the organism |
| Protocols | Different protocols for working with the organism, such as those for transformation |
| Biobrick parts | Biobrick parts such as promoters, reporters, etc designed to work in the organism |
| Vectors | Vectors that can be used for expression in this host organism |

## Supporting tools

### A. Codon Usage Table Generator (CUTE)

When heterologous genes are expressed in a host organism, codon optimization is routinely carried out for the heterologous gene to match the new host organism’s codon bias. To codon optimize a gene, it is required to know the codon usage table of the host organism. Kazusa(8) is a database of codon usage tables and the last data source used is from 2007. However, genomes for several organisms are being sequenced and codon usage tables need to be updated accordingly. Thus, we set out to create a tool that enables researchers to create codon usage tables from an input set of open reading frames. The GUI (Graphical User Interface) tool is implemented using JavaScript running on a webpage and requires an input list of ORFs in FASTA format. An output codon usage table is generated and displayed. The algorithm used for calculating is detailed below:

**Fig. 2.**
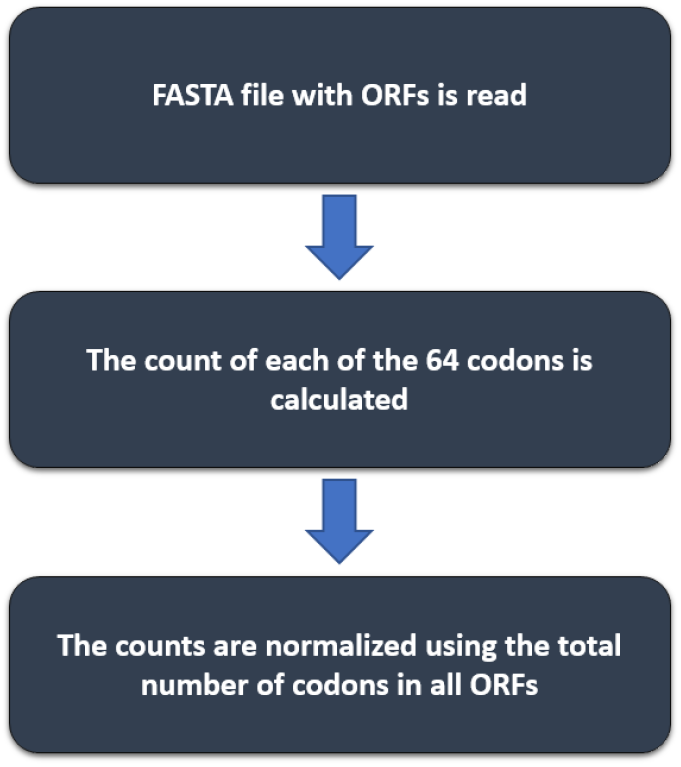
The algorithm used for Codon Usage Table Generator (CUTE)

The code is open source and available on GitHub, and the tool can be accessed at cute.chassidex.org

### B. Codon optimizer (COPTER)

We have also designed an interactive codon optimizer to optimize genes for expression in host organisms listed on ChassiDex. The tool is accessible at copter.chassidex.org.

**Fig. 3.**
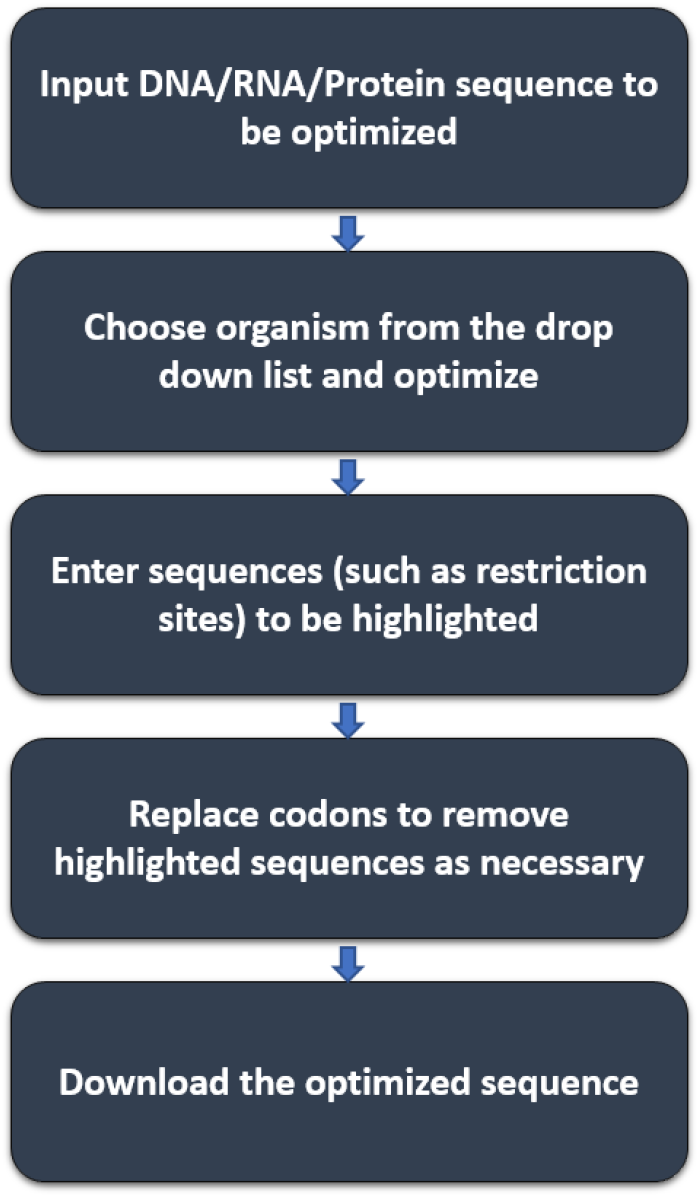
Workflow for using the Codon Optimizer (COPTER) tool

## Future Directions

- We hope to create a dedicated group to handle the curation of data into ChassiDex. The work will involve contacting iGEM teams and lab groups for data regarding the microorganisms.
- Establishing an online strain repository of the nonconventional microorganisms. This will be done by involving other labs willing to share material (strains, vectors and parts) under the Open Material Transfer Agreement (https://biobricks.org/openmta/). ChassiDex will act as an interface for the groups who wish to work with the chassis and the groups which can provide the chassis.
- Developing additional support tools that can aid the process of data mining across different data repositories and literature sources.

## Acknowledgements

We would like to thank the entire association of International Genetically Engineered Machine competition (igem.org) for giving us the opportunity to work on this database as our project for the year 2017 & 2018. We would like to thank Dr. Nitish Mahapatra at the Department of Biotechnology, Indian Institute of Technology, Madras who has been guiding this initiative since 2017 with his invaluable support as a primary faculty advisor. The entire team and associated mentors of team iGEM IIT Madras 2017 is credited for their valuable contribution of helping in the process of data mining and curation in the early stages of this database. iGEM teams from across the world (see Supplementary S1) have contributed to this growing database by adding and curating the data. We would like to thank the Bhupat and Jyoti Mehta School of Biosciences, Indian Institute of Technology, Madras for giving us the resources for this project. Lastly, we would like to thank the Indian Institute of Technology, Madras for providing us with the opportunity, funding and computational resources to build ChassiDex.

