## Supplemental file S2 for "ChassiDex: A microbial database useful for synthetic biology applications"

Supplementary data S2: Existing data curated by other iGEM teams


1) Pichia pastoris - curated by iGEM 2017 teams IISc-Bangalore and INSA-UPS France
2) Pseudomonas fluorescens - curated by iGEM 2018 Team SKLMT-China
3) Pichia pastoris SMD1168H and Vibrio harveyi JMH626- curated by iGEM 2017 team INSA-UPS France
4) Vibrio natriegens -  curated by 2018 iGEM team Marburg


For more information regarding collaborative efforts to curate and add data for aforementioned organisms in ChassiDex, please see the following links to the team wiki page:
1) http://2017.igem.org/Teams/Collaborations#IIT-Madras
2) http://2018.igem.org/Teams/Collaborations#collab_22
